# Comparison of eDNA, bulk-sample metabarcoding, and morphological approaches: A case study of riverine benthic macroinvertebrate communities

**DOI:** 10.1101/2023.05.30.542510

**Authors:** Arnelyn D. Doloiras-Laraño, Sakiko Yaegashi, Joeselle M. Serrana, Naoto Ishitani, Kozo Watanabe

**Author notes:** Corresponding Author: Prof. Kozo Watanabe, Ph.D.; Contact information; Center for Marine Environmental Studies (CMES), Ehime University, Bunkyo-cho 3, Matsuyama, Ehime 790-8577, Japan.

## Abstract

Freshwater biomonitoring is essential for aquatic biodiversity conservation. Advances in high-throughput sequencing allowed parallel sequencing of community samples containing DNA from environmental samples, i.e., metabarcoding. Two approaches of DNA-based method are widely used, bulk-sample metabarcoding the use of bulk tissues such as insects and eDNA the use of environmental samples such as air, water and soil. Despite the novelty of this approach for routine freshwater biomonitoring, questions still need to be answered about its applicability and reliability due to confounding factors, e.g., sample type, laboratory technicalities, and limitations of databases. Hence, studies on direct comparisons are essential to validate the efficiency of these molecular approaches compared to the conventional morphological approach to accurately assessed biodiversity for riverine benthic macroinvertebrate biomonitoring. This study used three approaches to estimate diversity and composition of benthic macroinvertebrates. We also evaluated the relationship between benthic macroinvertebrate communities and environmental factors. We morphologically identified 8,052 individuals from 35 families, 31 genera, and 29 species. eDNA metabarcoding identified 51 families, 84 genera, and 90 species, while 37 families, 55 genera, and 107 species were detected through bulk-sample metabarcoding. We report that bulk-sample metabarcoding showed finer taxonomic resolution than other approaches. Our study highlights the use of bulk-sample metabarcoding for macroinvertebrate biodiversity assessment.

## Introduction

The effects of global changes highlight the need for accurate data on species composition and effects of environmental factors such as flow discharge, temperature (Juergens et al., 2022; Jorgensen et al., 2010). Aquatic monitoring is not only essential to freshwater ecosystem but also to other water bodies ecosystem, because these ecosystems are directly affected by environmental factors such as water velocity, pH and temperature. Freshwater biodiversity represents a valuable natural resource that serves as a habitat for approximately 6% of all described species (Dudgeon et al., 2006). Freshwater systems across the world face serious ecological threats, making their protection and preservation crucial (Keck et al., 2017). Species loss compromises ecosystem resilience (Dudgeon et al., 2006); hence, freshwater biomonitoring remains one of the top priorities in ecological conservation. Here in this study, we utilized benthic macroinvertebrates as a case study to accurately measure the biodiverrsity. Benthic macroinvertebrates are conventionally used because of their sensitivity to water quality changes (Birk et al., 2012), river connectivity (Chiuh et al., 2020; Serrana et al., 2019) and flow regime (Miyake et al., 2021). The composition of benthic macroinvertebrates provides insights into changes in freshwater functions (Cao et al., 2018).

The traditional morphological approach is commonly used for biodiversity assessment in freshwater ecosystems. However, this conventional approach is time-consuming and labor-intensive (Pawlowski et al., 2018). Advances in DNA sequencing technologies have revolutionized biomonitoring by providing accurate data on species richness (Keck et al., 2017), and these drastically transformed how biodiversity surveys are conducted (Deiner et al., 2017). These offer in-depth insights and have become a potent tool for biomonitoring. Among these advanced technologies, DNA metabarcoding is a rapid, cost-effective, and practical approach to biomonitoring because it is rapid and inexpensive (Keck et al., 2017), producing massive amounts of data in a short time. This technique offers enhanced better estimates in biomonitoring data, ultimately transforming how species composition is determined (Taberlet et al., 2012). DNA metabarcoding routinely utilizes bulk samples and environmental DNA (eDNA) utilizes samples from water, soil, or sediments. While both employ almost the same techniques, sources of ideal samples and approaches specifically extracting species biodiversity data still need to be thoroughly explored.

Several studies compared two freshwater biomonitoring approaches at a time, i.e., morphological approach versus eDNA metabarcoding (Hajibabaei et al., 2020; Macher et al., 2018; Bratschen et al., 2017; Mächler et al., 2020; Gleason et al., 2021), or morphological approach versus eDNA metabarcoding (Blackman et al., 2017) and morphological approach versus bulk-sample metabarcoding (Serrana et al., 2019; Turunen et al., 2021; Emilson et al., 2017). Studies have found that all approaches have unique characteristics, advantages, and disadvantages because of confounding factors, i.e., nature of the sample, sampling method, laboratory, and bioinformatics analyses (Blackman et al., 2019; Barnes & Turner, 2016). Some of these studies do not apply both approaches to the same sample and have focused on a particular taxonomic group (Hajibabaei et al., 2012; Carew et al., 2013; Zhou et al., 2013; Gibson et al., 2014; Cowart et al., 2015; Zimmerman et al., 2015). Identification of benthic macroinvertebrates at the species level remains challenging. Incongruent results, such as the detection of target taxa and their relationship with environmental factors, impede their applications in biomonitoring, and the question of which method is the best to apply in biomonitoring needs to be answered (Aylagas et al., 2020). Thus, there is a need for thorough comparative analyses of the three approaches to test the effectiveness and reliability of macroinvertebrate biodiversity assessment for biomonitoring.

In this study, we compared eDNA metabarcoding against bulk-sample metabarcoding and morphological approach from the collected benthic macroinvertebrates in Shimanto and Niyodo Rivers in Shikoku Island, Japan. With this, we formulated four hypotheses; (1) we expect to observe higher taxonomic richness in bulk-sample metabarcoding compared to the morphological approach, which can be attributed to the identification of juvenile and damaged macroinvertebrate samples that are usually cannot identify during the morphological identification. Similarly, a higher taxonomic richness from bulk-sample metabarcoding than in eDNA metabarcoding is expected to be detected because it targets the whole organism. In contrast, eDNA metabarcoding targets traces of DNA expelled into the environment. In addition, we expect to observe finer taxonomic resolution from bulk-sample metabarcoding because of its ability to detect taxa at the species level compared to morpgological appproach. (2) We expect higher beta diversity in bulk-sample compared to eDNA metabarcoding because each sampling site was different in terms of environmental factors. (3) We expect a positive correlation between sequencing reads, morpho-taxa and abundance. (4) Finally, we expect a stronger association of the environmental factors to community composition from bulk-sample metabarcoding due to its better representation of the local environment.

## Methods

### Study site and sample collection

The field surveys were conducted between July and October 2016 in Shimanto and Niyodo Rivers, located on Shikoku Island, Japan (Figure 1). The Shimanto River comes from its headwater on the Mount Irazu slopes and flows into Tosa bay. The total length of the Shimanto River is 196 km, and the river has a basin area of approximately 2,270 km^2^. The Niyodo River originates from its headwater on Mount Ishizuchi with 1,982 meters high slopes and flows 124 km. The Niyodo River covers an approximate area of 1,560 km^2^ and is surrounded by 1,287 ha of vegetation and forest (MFAJ, 2019). The Niyodo River is known for the best water quality in Japan, and the Shimanto is a continuous river with few dams in its watershed.

**Figure 1.**
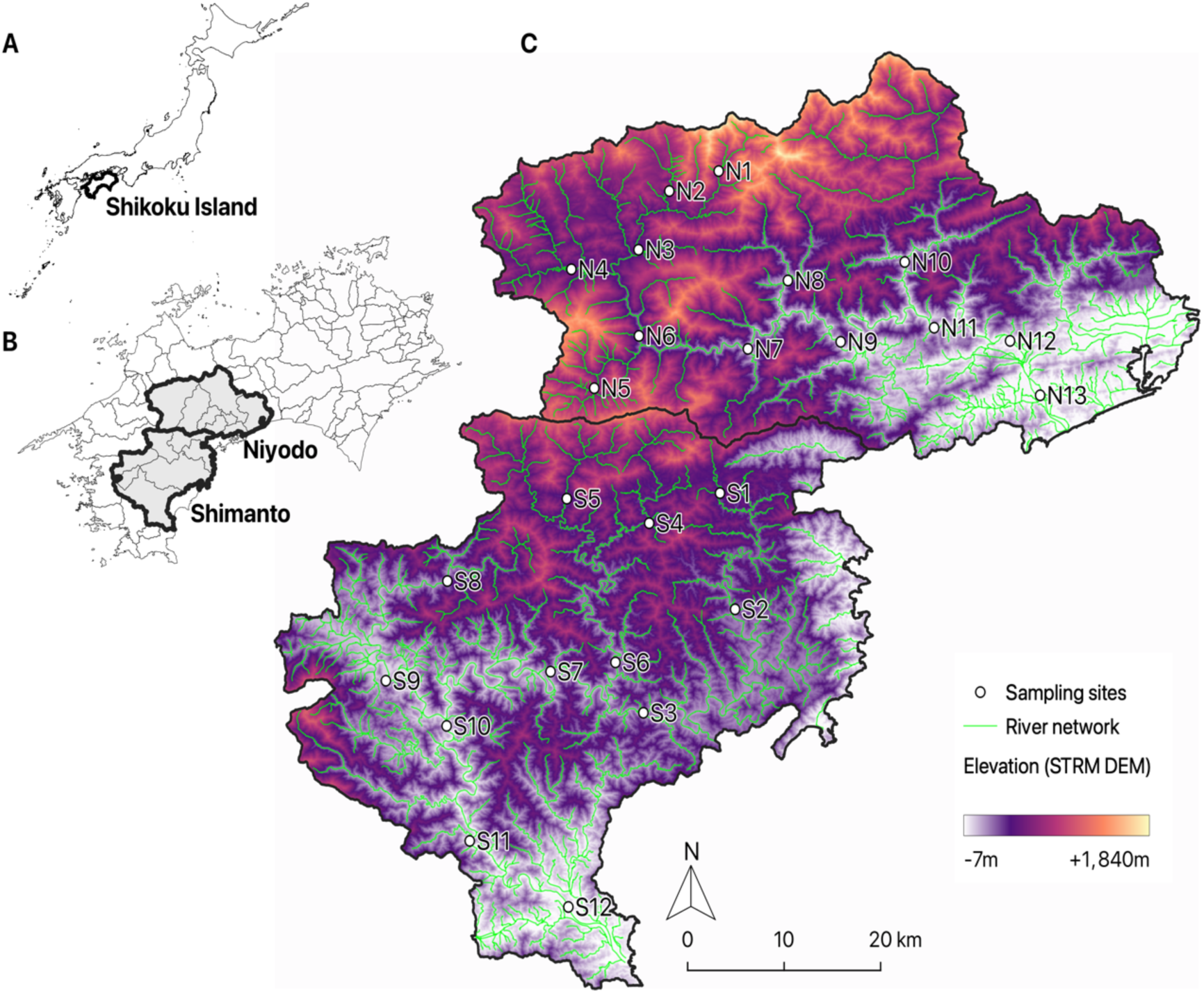
Map of the sampling sites. (A) shows the location of Shikoku Prefecture in Japan; (B) Shows the location of Niyodo and Shimanto Rivers on Shikoku, Island. (C) Shows the 13 sampling sites (N1 to N13) in Niyodo River and 12 sampling sites (S1 to S12) in Shimanto River.

Water samples were collected from Shimanto River (n = 12) and Niyodo River (n = 13) with two replicates (n = 50). Sampling sites are located approximately every 10-30 km from upstream to downstream of the mainstream to cover the various segments (up-, mid- and downstream) within the basins. As tributary reaches can have different habitats than the mainstream, 1-2 additional sampling sites were assigned to major tributary area. We conducted the samplings under ordinaly water discharge. Each site had a different physical environment (i.e., water depth, water current velocity, river sediment), water quality (i.e., pH, DO, dissolved nitrogen and phosphorous) and potentially food resources of aquatic insects (i.e., Periphyton biomass). Ten liters of surface water from the mainstream was collected in each sterile water bag. A biological blank as a negative control was used by filtering one liter of ultra-pure water. The 1.0 L of surface water samples were vacuum-filtered through a 45 mm MCE membrane filter (Advantec, 0.22 µm pore size) and stored at -20°C before processing for molecular analysis.

Macroinvertebrates were collected using a Surber net (25 x 25 cm quadrat, 0.5 mm mesh) with three replications (n=75) at the same locations and time as the water sampling. The Surber net with a 25 cm x 25 cm quadrat frame was placed on the river bed, and benthic animals were collected by brushing the stones and gravels and by kicking the river bed within the frame. Sample collections were carried out three different location on rapid area within the same reach in a sampling site. Surber samples were preserved in 99.5% ethanol in the field, transported to the laboratory, rinsed carefully to remove the sediments and other organic matters (i.e., leaves, wood braches) without losing any macroinvertebrates, transferred to new 99.5% ethanol in order to preserve DNA stably for long time (Stein et al. 2013, Marquina et al. 2021), and stored at 4 °C before processing for morphology-based identification and later DNA analysis.

Environmental factors were collected from each study site at the same time when sampling was carried out in order to characterize the relationship with macroinvertebrate composition (Table 1). We mesured on three types of environmental factors; phisical habitat structure (i.e., current velocity, water depth, water temparature, river sediments), water quality (i.e., pH, dissolved oxygen, electric conductivity) and potentially food resources for aquatic insects (Chlorophyll *a*) on each site (n = 25). Water current velocity and water depth were measured simultaneously at ten randomly different locations. A portable current meter flow probe (Global Water, USA) measured water current velocity at half of the depth of the water. We employed the average values of water depth and velocity for environmental analysis. Water temperature, dissolved oxygen (DO), water pH, and electric conductivity (EC) were measured using a portable water quality meter (DO-31P, TOADKK) for the water temperature and the dissolved oxygen, Laqua twin pH-11B (Horiba) for the pH, EC-33B (Horiba) for the EC, respectively. Surface water samples were collected from each site for water chemistry analysis. Chlorophyll *a* concentration was measured to estimate Periphyton biomass, which is one of food resources for macroinvertebrates, in the range of 5 cm^2^. Periphyton was scrubbed off a brush, and Chlorophyll was extracted with 99.5% ethanol. The extracts were measured by a spectrophotometer (Model u-1100; Hitachi Co., Tokyo, Japan), and Chlorophyll *a* concentration was calculated based on Unesco,1969. Collected water samples measured for water quality analysis. Ammonium nitrogen (NH_4_-N), nitrate-nitrogen (NO_3_-N), nitrite-nitrogen (NO_2_-N), and Phosphorous (PO_4_-P) were measured using Auto Analyzer 3 (BRAN-LUEBBE). The components of river sediments also examined because some benthic animals use sand, mud and gravel as their habitats. We were collected 1-3 kg of river sediments including gravels, sand and silt from a sandbar adjacent to the water channel once in each site and assorted by five different sieves (mesh size: 0.75 mm, 0.25 mm, 0.85 mm, 2 mm, 4 mm) after drying them. Grain size accumulation curves were described by the assorted sediments weight (i.e., more than 4mm (middle or larger gravels), 2mm (small gravels), 0.85mm (large sand), 0.25mm (middle sand), 0.75mm (small sand), thess than 0.75mm (silt)). Finlally we calculated sediment size of 10%wt (D10), 30%wt (D30), 50% wt (D50), 60% wt (D60), and uniformity coefficient Uc (= D60/D10).

### Morphological analysis

Macroinvertebrates samples were sorted and taxonomically identified based on the keys to the aquatic insects of Japan by Kawai and Tanida (2005) using a stereomicroscope (SZ-61, OLYMPUS). The abundance (number of individuals) per sampling site was counted. After the sorting and identification, total dry biomass (mg) was measured using UMX2 Ultra-microbalance (Mettler-Toledo., USA). The macroinvertebrate samples were then reconstituted, preserved in 99.9% ethanol, and stored at 4°C for molecular processing.

### DNA extraction, library preparation, and sequencing

Dried macroinvertebrates from each sampling site (n=25) were then homogenized into a 1.5 ml tube. The lysis buffer composed of 300 µl of PBS,180 µl of ATL buffer (Qiagen), and 20 µl of proteinase K solution (Qiagen) were added into the tube. It was incubated overnight at 56°C in a water bath. Next, 500 µl of PCI (Phenol: Chloroform: Isoamyl alcohol = 25:24:1) was added and mixed well. The tube was centrifuged (12000 rpm, 10 minutes, room temperature), and supernatant was transferred to a new 1.5 ml tube. Then, 100 µl of chloroform was added, mixed well, and centrifuged under the same condition. The supernatant was transferred to a new 1.5 ml tube, and 30 µl of 3M sodium acetate (pH 5.2) and 660 µl of 99.5% ethanol were added. After the mixture was stored at -20°C for 2-3 hours, it was centrifuged (12000 rpm, 15 minutes, room temperature). After the supernatant was discarded, 1 ml of 70% ethanol was added and centrifuged (12000 rpm, 15 minutes at room temperature). The supernatant was discarded again, and the pellet was dried. Finally, the DNA was resuspended using 50 µl of TE buffer.

Environmental DNA was extracted by the phenol-chloroform extraction method. First, each filter was cut by sterilized scissors and put in a 50 ml tube with 10 ml of HMW buffer (final concentration; 0.1M Tris, 0.1M EDTA, 0.7M NaCl), 6.5 µl of proteinase K (Qiagen), and 6.5 µl of 10% SDS. The tube was incubated for 2 hours at 55 °C in the water bath. Next, 650 µl of TE-saturated phenol was added and mixed well. The tube was centrifuged (10000 x g, 10 minutes, room temperature), and the supernatant was transferred to a new 50 ml tube. Then, 225 µl of TE-saturated phenol and CIA (Chloroform: Isoamyl alcohol = 24:1) were mixed well into the tube. We performed centrifugation and transferred the supernatant to a new tube again. Then, 100 µl of 3M sodium acetate (pH 5.2) and 20 ml of 99.5% ethanol were mixed and preserved at -20 °C for 1 hour. The mixture was centrifuged (10000 x g, 20 min, 0–4 °C), and the ethanol was discarded. Then, 20 ml of 70% ethanol was added into the tube and centrifuged in the same condition. The 70% ethanol was discarded, and the tube was dried. The DNA was resuspended using 500 µl of a TE buffer (pH 8.0). Finally, extracted DNA was purified using the One-Step PCR Inhibitor Removal Kit (Zymo Research) and was stored at -20 °C.

Purified genomic DNA was quantified on a NanoDrop spectrophotometer (Thermo Scientific, USA. Nanodrop 2000). Ten times diluted eDNA and 100 times diluted bulk DNA were used for two-step PCR. The first PCR step (35 cycles) was conducted using the primer set BF2 (GCH-CCH-GAY-ATR-GCH-TTY-CC) + BR2 (TCD-GGR-TGN-CCR-AAR-AAY-CA) of Elbrecht and Leese (2017). PCR reaction solution consisted of 3 µL of 5X Phusion HF Reaction Buffer (New England Biolad), 0.6 µL of 2.5 mM dNTPs (TaKaRa, Japan), 0.15 µL of Phusion HF DNA Polymerase (2000U/ml) (New England Biolabs), 0.45 µl of 50 mM MgCl2 Solution (New England Biolabs), 0.45 µl of DMSO (New England Biolabs), 0.75 µl of 10 mM each BF2 and BR2 primer, 0.5 µlof each diluted each DNA. The total volume was adjusted to 15 µL with PCR-grade water. The first PCR condition was as follows: initial denaturation at 94 °C for 3 mins; 35 amplification cycles of 94 °C for the 30s, 58 °C for 30s, and 72 °C for 1 min; and final extension at 72 °C for 10 mins, and stored at 4°C.

The second PCR step was done to add the adaptors and barcodes. We prepared a PCR reaction mixture consisting of 3 µL of 5X Phusion HF Reaction Buffer (New England Biolad), 0.6 µL of 2.5 mM dNTPs (TaKaRa), 0.25 µL of Phusion HF DNA Polymerase (2000U/ml) (New England Biolabs), 0.6µl of 50 mM MgCl2 Solution (New England Biolabs), 0.45 µl of DMSO (New England Biolabs), 0.75 µl of 10 mM each forward and reverse primer, 0.5 µl of 1st PCR products. The total volume was adjusted to 15 µL with PCR-grade water. For Illumina sequencing system, we designed the 2nd PCR primer following the pattern P5/P7 sequence+sequence primer region+Tag sequences (6bp) +BF2/BR2 primer sequences (Table S1). The temperature conditions were as follows: heating at 94°C for 2 min, 15 cycles of following steps by thermal denaturation at 94°C for 30 s, annealing at 55°C for 30 s, and extension reaction at 72°C for 1 min. Finally, an extension reaction was performed at 72°C for 10 min and stored at 4°C. T100 Thermal Cycler (Bio-RAD) was used for both PCR. PCR products were then visualized on 1.5% agarose gel to confirm clear bands for successful PCR amplification. The amplicon size was 421 bp. Negative control was used to monitor contamination from amplicon library preparation, and no quantifiable information amplicon was detected for further statistical analyses.

A total of 127 amplicon libraries were constructed, i.e., one negative control, duplicates of 25 sites for eDNA, and triplicates of 25 sites for bulk-sample. Next, we quantified each PCR product using Kapa Library Quantification Kit (Kapa Biosystems, Massachusetts, USA) in C1000 Touch, Real-Time System Thermal cycler (Bio-Rad Laboratories, CA, USA) and normalized it to 100 nM concentration. The final library was constructed by pooling an equimolar concentration of each amplicon library. The library was purified using SPRI Select (Beckman Coulter Inc., USA), following the manufacturer’s protocol. We used the Agilent Bioanalyzer 2100 system and Agilent High Sensitivity DNA kit (Agilent Biotechnologies) to assess the quality of pooled library by obtaining a single clear band of 605 bp. Sequencing was conducted on an Illumina Miseq using the Miseq Reagent Kit v3, 300 cycles (Miseq, Illumina, USA, MS-102-3003) (paired-end 2x 300 bp) with a starting concentration of 4 nM, denatured to a final concentration of 6 pM and used 20% concentration of Phix (Miseq, Illumina, USA).

### Read processing and taxonomic assignment

Raw paired-end reads were quality-checked using FastQC v.0.11.8 (Andrews, 2010). The raw reads were demultiplexed via JAMP v.0.67 (http://github.com/vascoElbrecht/JAMP) (Elbrecht et al., 2018) in R v.4.0. After demultiplexing, primer sequences were trimmed using Cutadapt v.2.1(Martin, 2011). Then, the software package DADA2 v.1.12 was used to infer amplicon sequence variants (ASVs) using the methods from Callahan et al. (2016) in R v.4.0. ASVs were concluded after trimming, quality filtering, denoising, merging, chimera and singleton removal. Minimum length of 220 bp and maximum error expected (--maxee) 3 and 5 for forward and reverse reads, respectively were used as parameters in quality filtering.

ASV sequences were subjected to Barcode of Life Databases (BOLD) searches wherein the top hit with a sequence identity of ≥97 %, for species assignment of each representative sequence using the Boldigger (Buchner & Leese, 2020). The morphology-based identification approach identified macroinvertebrates into broad and inconsistent taxonomic levels, while the DNA metabarcoding approach identified most macroinvertebrate samples up to the species level. The morphology-based identification, bulk-sample metabarcoding, and eDNA metabarcoding (morpho-taxa, bulk-sample meta-taxa, eDNA meta-taxa), the DNA metabarcoding species were collapsed into the taxonomic level in parallel to the morpho-taxa. Taxa that are unique for each method are kept as it is.

### Statistical analyses and visualization

Statistical analyses and visualization were done in R v.4.0 (R Core Team, 2016). Alpha diversity (species diversity within samples) and beta diversity (species diversity between samples) were calculated using the vegan package in R v.4.0 (Oksanen, 2022). Alpha diversity of the three datasets was evaluated using the alpha metrics; Chao 1 richness (Chao, 1984), Simpson’s diversity index (Simpson, 1949), and the Shannon-Weaver diversity index (Shannon & Weaver, 1949). The shared taxonomic profile based on richness was visualized using Venny (Oliveros, 2015).

Beta diversity was calculated using the vegan package v.2.6-4 (Oksanen,2022). Non-multidimensional scaling (NMDS) analysis and permutation of ANOVA (PERMANOVA) were done to test the significant difference between the bulk-sample metabarcoding and eDNA metabarcoding approaches. We also analyzed similarities (ANOSIM) to test whether there was a statistically significant similarity between the two metabarcoding approaches of sampling units. We tested the differences between bulk-sample metabarcoding and eDNA metabarcoding (permutation = 9999, alpha = 0.05). All statistical analyses were performed in the vegan package of R v.4.0 (R Core Team, 2016). Distance-based redundancy analysis (dbRDA) was performed in vegan package in R v.4.0 (Legendre & Gallagher, 2001) to evaluate the relationship between environmental factors and macroinvertebrates in three approaches.

To further evaluate the correlation between sequence reads, abundance, and biomass, we plotted the number of sequence reads of given macroinvertebrate taxa in DNA metabarcoding sequencing reads against the total quantity and total biomass of the morphology-based identification. Pearson’s correlation analysis on log-transformed values was also performed. The spatial scale of bulk-sample and eDNA metabarcoding were determined using spatial autocorrelation using the Bray-Curtis dissimilarity index and the computed air and water geographic distance of the sampling sites. We calculated the autocorrelation coefficient (*r*) from the geographic distance and the genetic distance (beta diversity) using the GenAlex program. Mantel tests for air and water distance were done using a vegan package in Rstudio v.4.0 (R Core Team, 2016).

## Results

### Taxonomic identification

A total of 8,052 individuals were collected from 25 sampling sites and morphologically identified (Table 2). Based on our expertise, we have morphologically identified 69 taxa to the finest taxonomic rank possible. These were distributed across 35 families, 31 genera, and 29 species. The most abundant order was Ephemetroptera (4,650 individuals, 57.8%), followed by Trichoptera (1,485, 18.4%), Diptera (1,302, 16.2%), Plecoptera (414, 5.1%), Coleoptera (186, 2.3%), Odonata (12, 0.2%), Amphipoda (2, 0.02%) and Isopoda (1, 0.01%). The top three dominant species were *Uracanthella punctisetae* (1,133 individuals,14.1%), *Epeorus latyfolium* (415,5.2%) and *Paraleptophlebia chocolate* (329, 4.1%) (Table 2).

A total of 12,184,167 raw reads were generated after Miseq sequencing. After the quality filtering, 8,682,259 sequencing reads were retained. The bulk-sample dataset generated 710,375 while the eDNA dataset generated only 11,931 sequencing reads (Table S1). A total of 7,456 ASVs were generated, with 796 ASVs (10.68%) assigned to taxa with 97% similarity with BOLDigger (Buchner and Leese, 2020). Then, a total of 632 taxonomically assigned-ASVs were inferred from the bulk-sample dataset, and 164 ASVs for the eDNA metabarcoding dataset. eDNA metabarcoding identified 51 families, 84 genera, and 90 species, while 37 families, 55 genera, and 107 species were detected through bulk-sample metabarcoding.

### Comparison of morphological approach, bulk-sample, and eDNA metabarcoding

Alpha diversity was highest in bulk-sample metabarcoding than in morphological approach and eDNA metabarcoding (Figure 2A). The NMDS ordination showed a clear difference between the three datasets i.e. morphological approach, bulk-sample and eDNA metabarcoding datasets (Figure 2B). The PERMANOVA (r^2^ = 0.2546, *p* = 0.001) showed a statistically significant difference between the bulk-sample metabarcoding and eDNA metabarcoding approaches. Results also showed that communities significantly differed between the bulk-sample metabarcoding and eDNA metabarcoding approaches using ANOSIM (R = 0.563, *p* = 0.001).

**Figure 2.**
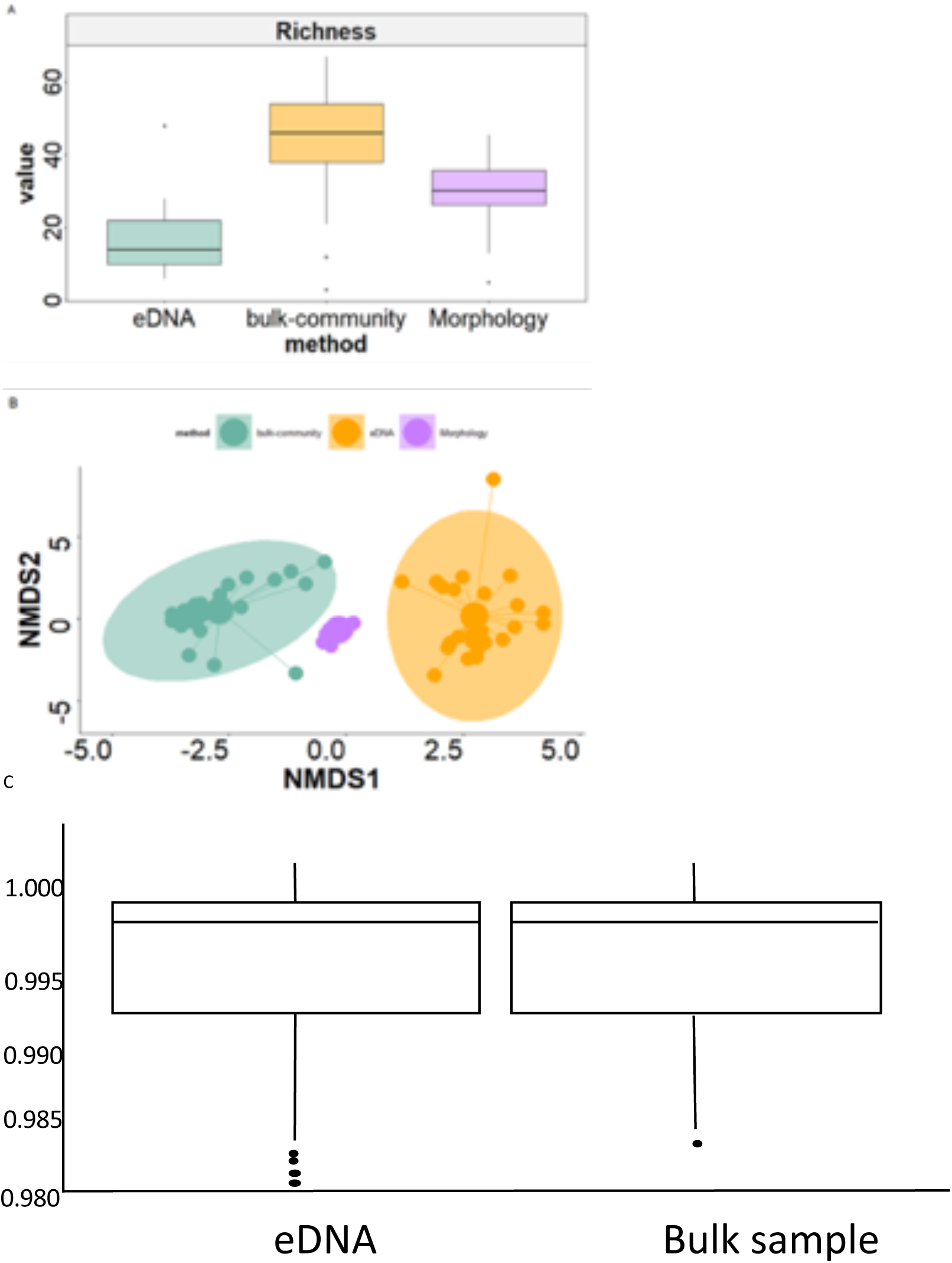
(A) The taxa richness per approach .e morphology-based identification and bulk-sample metabarcoding and eDNA metabarcoding; (B) Non-multidimensional scaling of three approaches show the clustering and separation; (C) Homogeneity test of bulk community and eDNA metabarcoding.

### Correlation analyses

A significant positive correlation was found (R = 0.27, *p* = 0.025) between the read abundance of meta-taxa and the total abundance of morpho-taxa (Figure 3A). Also, a positive correlation was detected between the read abundance of meta-taxa and the total biomass of morpho-taxa (R = 0.4, *p* = 0.0065) (Figure 3B). No significant correlation was detected in spatial autocorrelation between the geographic distance and beta diversity (Figure S4).

**Figure 3.**
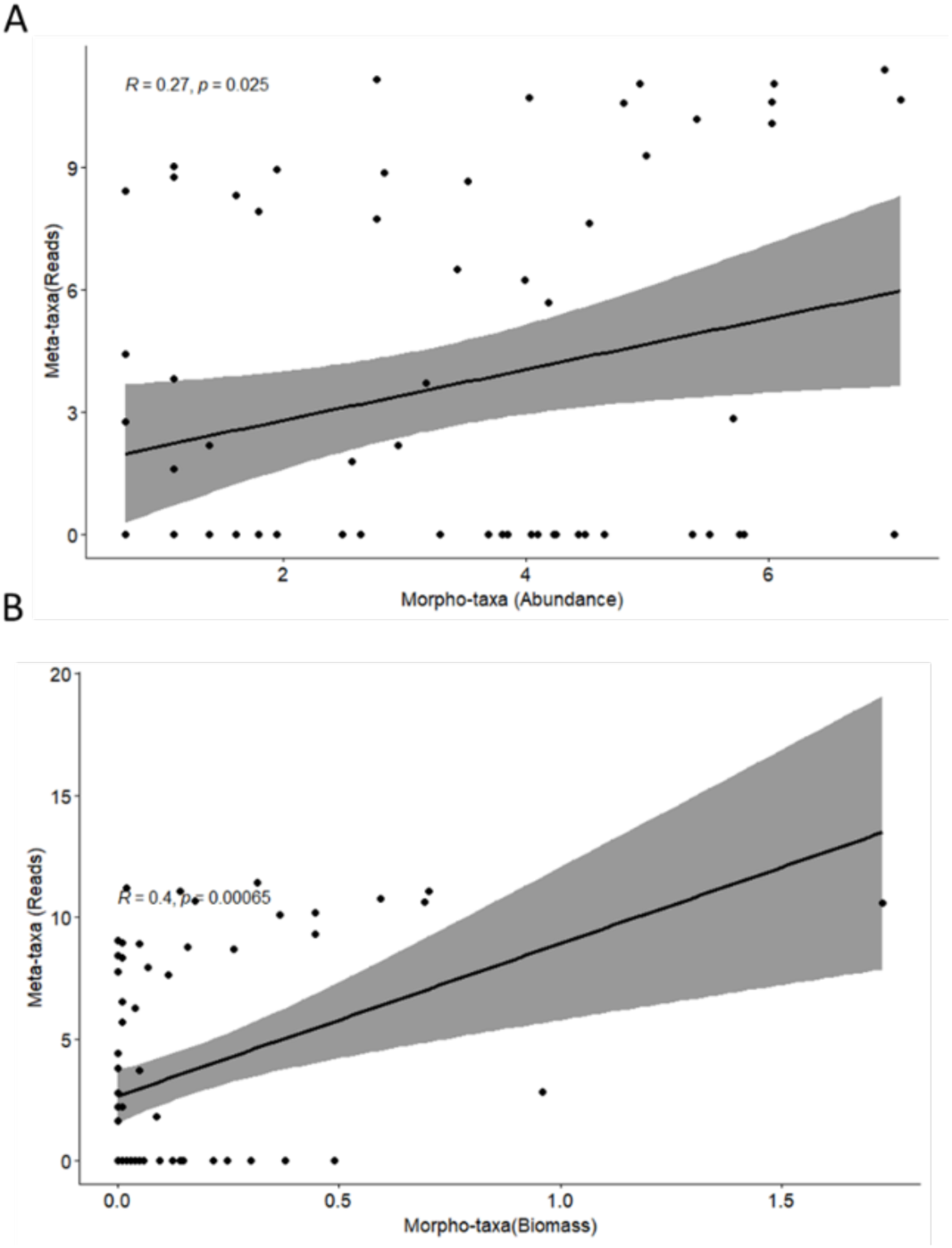
Correlation between (A) Morpho-taxa (Abundance) and meta-taxa (Sequence Reads) (B)Morpho-taxa (Biomass) and meta-taxa (Sequence Reads).

### Environmental parameters and macroinvertebrate communities

The dbRDA results showed that three physicochemical parameters, i.e., water velocity (p=0.001), water depth (*p* = 0.011), and sediment influenced the bulk-sample metabarcoding approach (Figure 4A & 4B). NH_4_ (*p* = 0.0321), was selected in the morphology-based identification approach. However, no physicochemical parameter was found in eDNA metabarcoding approach (Figure 4C).

**Figure 4.**
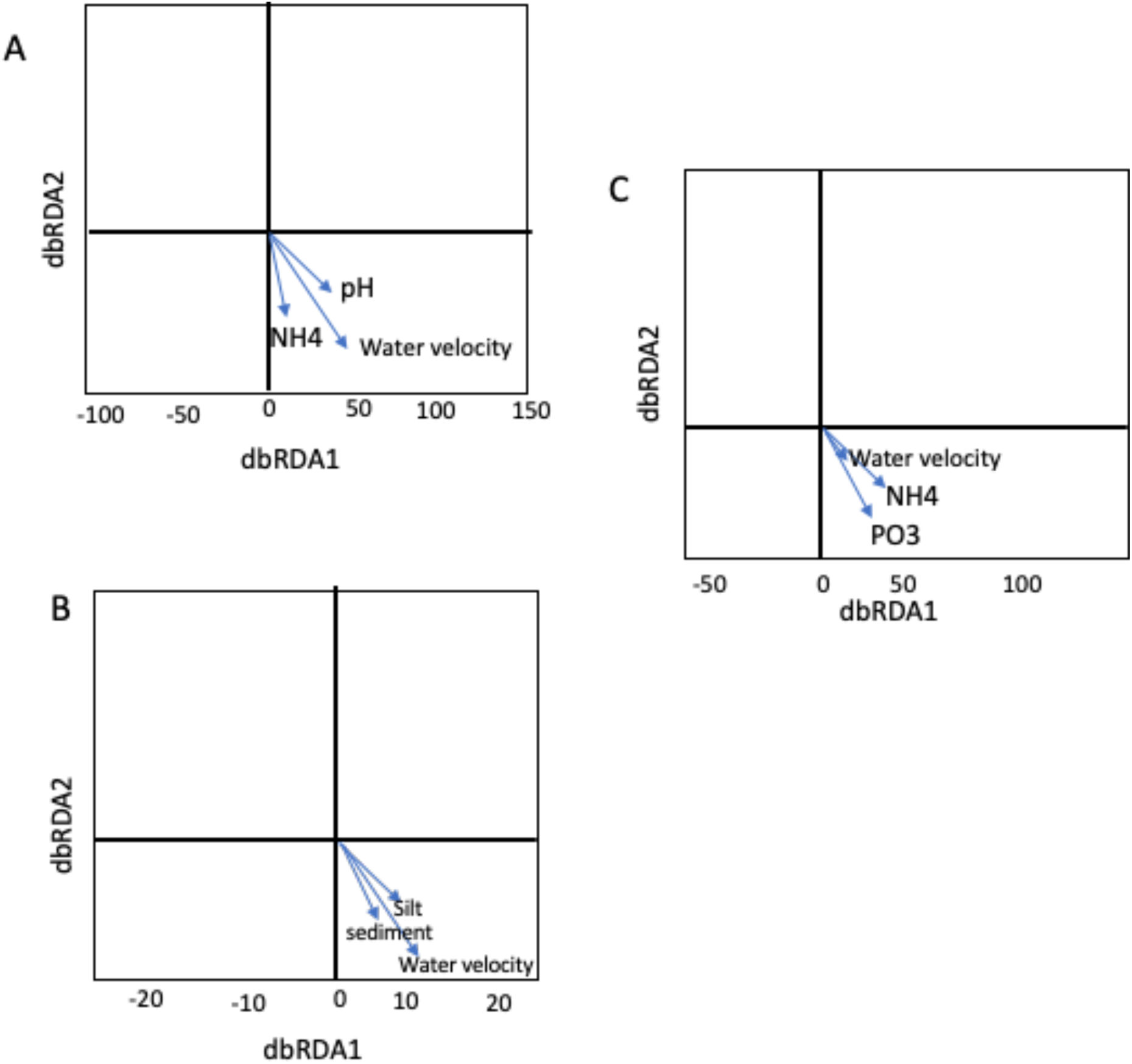
Distance-based redundancy analysis of three approaches; (A) bulk-sample metabarcoding, (B) morphology-based identification approach, (C) eDNA metabarcoding.

**Figure 5.**
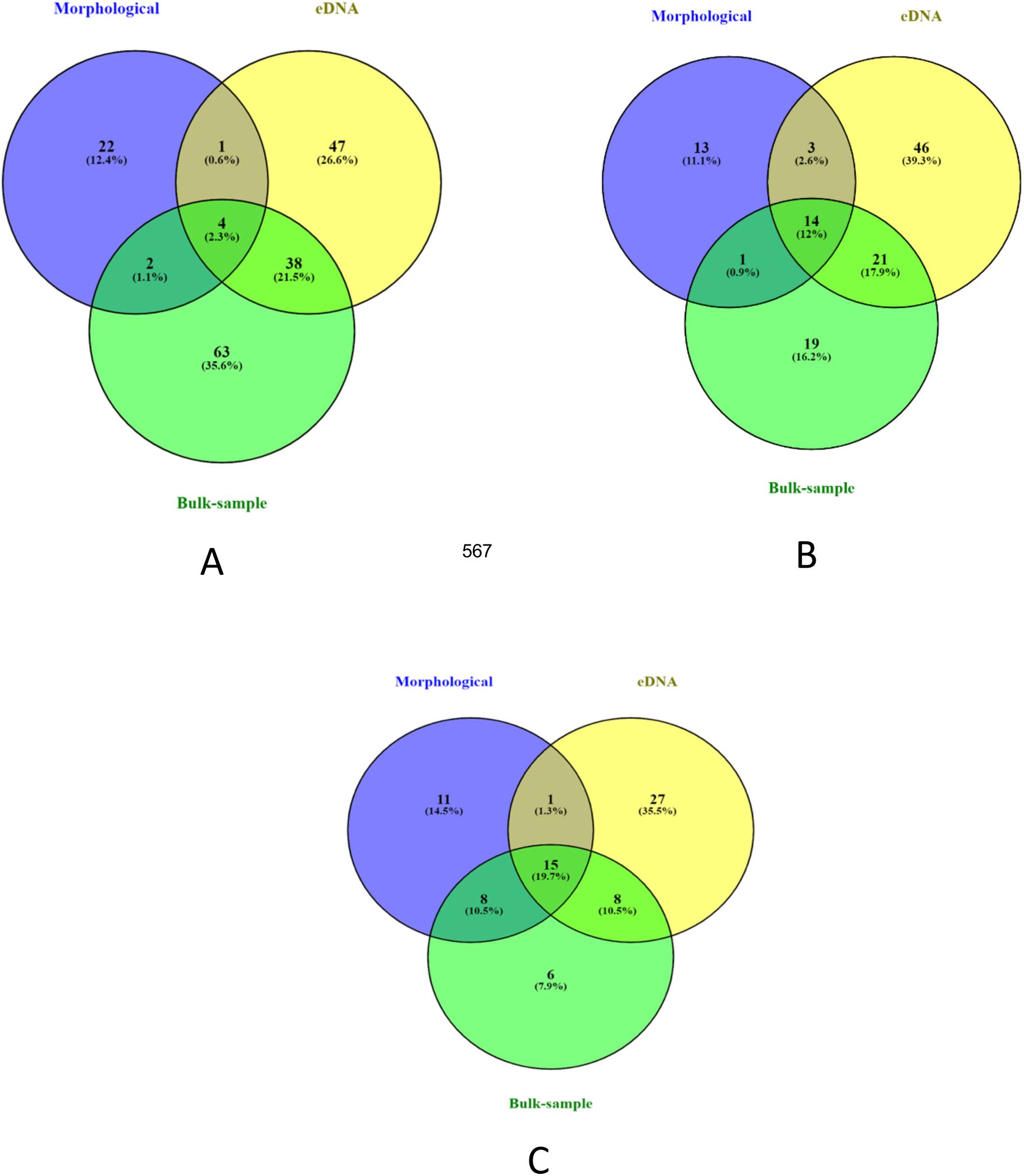
Venn diagrams of taxa sharing by the three approaches, e.g., morphological analysis approach, bulk-sample metabarcoding, and eDNA metabarcoding based on the collapsed table (A) Species Level; (B) Genus Level; (C) Family Level.

## Discussion

### Comparison of bulk-sample metabarcoding and morphological approach

Our study showed differences between bulksample and morphological approach. Bulk-sample metabarcoding also provided a finer taxonomic resolution at the species level (n=107). Taxonomic resolution is the ability to identify taxa to finer taxonomic levels such as the species level. Our results were in congruence with previous benthic macroinvertebrates studies (Elbrecht et al., 2017; Hajibabaei et al., 2019; Gleason, 2020), which showed the capability of the bulk-sample metabarcoding approach to identify taxon diversity in complex groups, and samples in larval stages in which identification is limited due to the absence of defining morphological characters, e.g., Diptera (Chironomidae and Simuliidae) and Trichoptera (Limnephilidae) (Hajibabaei et al., 2019).

False-negative results were shown when bulk-sample metabarcoding revealed 41 more species not detected when the morphological approach was employed. This result is expected since the morphological method is unsuitable for identifying early-stage macroinvertebrate and damaged samples since not all taxonomic characters required for identification can be observed in the samples. As a result, the classification combines them into higher taxonomic groups such as family or order. Since the taxonomic resolution is confined to higher taxa, only limited information is available to assess benthic macroinvertebrate biodiversity accurately. This point to the advantage of using DNA sequences from bulk-sample metabarcoding. The capability of bulk-sample metabarcoding to identify damaged and juvenile specimens overcomes the challenges in morphological analysis. These results are consistent with previous studies of macroinvertebrate diversity in aquatic ecosystems (Turunen et al., 2022; Serrana et al., 2019; Baird & Sweeney, 2011; Sweeney et al., 2011; Stein et al., 2013a; Gibson et al., 2015; Elbrecht et al., 2017).

Despite the advantages of bulk-sample metabarcoding over the morphological analysis approach, false-positive detection may also be a case in bulk-sample metabarcoding (Table 1). For example, uncommon or rare species may not be detected or underrepresented since abundant taxa can mask these species during molecular processing, such as DNA extraction, PCR amplification, and sequencing. The bioinformatics analyses may also be affected due to low or no sequence read abundance (Evans et al., 2016; Elbrecht et al., 2018; Schenk et al., 2019) or lack of deposited information in sequence reference databases. Furthermore, the detection of macroinvertebrates by bulk-sample metabarcoding can be strongly influenced by various laboratory methodologies, such as primer mismatches that prevent the amplification of specific taxonomic groups by PCR, undetected chimeras, and misidentified taxonomic groups (Deagle et al., 2014; Pinol et al., 2014; Elbrecht & Leese, 2015). Nevertheless, given the finer taxa resolution using bulk-sample metabarcoding and its capability to identify samples with incomplete taxonomic characters, this can still be a potent tool for species-level detection studies.

### Comparison between bulk-sample metabarcoding versus eDNA metabarcoding

Our results showed that bulk-sample metabarcoding could capture a higher number of identified benthic macroinvertebrates compared to eDNA metabarcoding, as this represents DNA extracted from the bulk-sample of benthic macroinvertebrates. In contrast, the DNA from the eDNA metabarcoding water samples tends to be diluted and low in concentration as these are just intracellular and extracellular DNA released from the benthic macroinvertebrates to the aquatic environment. Also, the DNA has different concentrations and greatly varies because DNA must be released in the aquatic environment and differs among macroinvertebrates. Our results highlight the effectivity of bulk-sample metabarcoding than eDNA metabarcoding, characterizing and representing the local macroinvertebrate communities in a given sampling site.

Moreover, DNA from benthic macroinvertebrates living in water, on the surface of stones, mud, and gravel, is difficult to detect. Given this ecology, it is expected that the DNA of benthic macroinvertebrates is only sometimes present in the water and will not accurately represent the local benthic macroinvertebrate communities in a given sampling site. eDNA metabarcoding can also cause a disparity in inferred biodiversity patterns when used in spatiotemporal analysis since the DNA present may have been transported from another site or just remnants or feces of a species that were active and not necessarily present at the specific location or time (Deiner et al., 2016). eDNA metabarcoding may be utilized in conducting multi-taxa diversity assessment using the same sampling method and DNA template (Campbell et al., 2022). The intrinsic nature of eDNA may differ between multicellular eukaryotes and prokaryotes or micro-eukaryotes. There is a higher probability of detecting intracellular DNA from microbes, as intact individuals are more readily isolated from lower DNA concentrations.

### Macroinvertebrate community-environment relationship

Our dbRDA analysis showed a stronger and clearer association between macroinvertebrate community structure, water velocity, and sediment in the bulk-sample metabarcoding approach. Water velocity is the driving force for delivering external nutrients to macroinvertebrates (Soballe & Kimmel, 1987). This correlation is evident between the higher number of identified taxa in the bulk-sample metabarcoding approach. Environmental changes can strongly impact benthic macroinvertebrate ecosystems (Dou et al., 2022; Mao et al., 2023).

The environmental NH_4_ was found to be correlated with the morphology-based identification approach. The community with those species grouped into one taxonomical level may reflect changes in the specific factors (e.g., NH_4_ in our study), respectively. The grouping has the averaging effect neutralizing the opposite responses and strengthening similar responses. No ecological factor was significantly correlated to eDNA metabarcoding, which may indicate that eDNA metabarcoding cannot represent the local environment composition and heterogeneity. Also, there may be a better approach for the environmental factor assessment of a specific sampling site of interest (Macher et al., 2018). This is because bulk sample was collected locally while the eDNA represents a composite sample from the upstream watershed. Therefore, thez have different origin. This implied that bulk sample correlated with the local environment factors and eDNA metabarcoding correlate with other factors more related to watershed features. However, at a global scale, eDNA can be a possible alternative to assess at a higher scale which can provide information on different environmental factors at regional levels (e.g., land use in upstream catchments), despite the difficulty in the collection. Overall, bulk-sample metabarcoding connects with more environmental factors and is an efficient approach for assessment, at least in the river system included in this study.

### Correlation analyses between approaches

Our results showed a significant positive correlation between bulk-sample sequence read abundance and morpho-taxa abundance. This result suggests the potential of bulk_sample metabarcoding for measuring macroinvertebrate communities based on the sequence read information. We also found a significant positive correlation between morpho-taxa biomass and sequencing read abundance. This finding supported our hypothesis that large macroinvertebrates with high biomass would be overrepresented. Our results implied that high biomass would be overrepresented, for the extraction of DNA material in bulk and smaller organisms lead to a higher number of sequence reads (Serrana et al., 2019; Elbrecht et al., 2017). This could be explained by large-bodied macroinvertebrates contributing to the total body mass are expected to have a higher sequence read abundance than small-bodied macroinvertebrates. This finding is the same as the observation in the experiment of Elbrecht and Leese (2015, 2017) that showed the biomass of smaller species does not lead to their systematic exclusion, provided that the specimens have similar amplification efficiencies. These suggest that upscaling with more complex macroinvertebrate taxa might be possible. However, the quantification of abundance depends on the reliable recovery of taxonomic abundance (Gotelli (& Chao, 2013; Aylagas et al., 2014; Bucklin et al., 2016). Previous studies on benthic macroinvertebrates showed a distinct positive correlation between biomass and sequence reads (Elbrecht & Leese, 2015; Serrana et al., 2021). Another study showed a positive correlation between the number of reads and species abundance (Yu et al., 2012) because highly abundant species will have higher concentrations of template DNA (Cowart et al., 2015; Machler et al., 2019).

## Conclusion

Our current study provided different results of the three approaches widely used in aquatic biomonitoring. We showed that each approach had its own advantage and disadvantage in aquatic biomonitoring. Hence. there still a need to standardize the methods such as DNA extraction, library preparation ad bioinformatics used to have better comparisons of the DNA-based approaches with the morphological. We recommend to further exploring eDNA metabarcoding by increasing the number of sampling effort that can be analyzed.

## Data availability

The raw sequence data is available through NCBI SRA under the BioProject number PRJN867110.

## Author Contribution

SY and KW conceptualized and designed the study; SY conducted the fieldwork and the morphological identification; NI performed the laboratory works; ADL and JMS performed the metabarcoding analyses. ADL, SY, JMS, and KW interpreted the results. ADL and KW drafted the manuscript with writing inputs and revisions from all authors.

## Acknowledgment

This research was financially supported by the Japan Society for the Promotion of Science (JSPS) Grant-in-Aid for Scientific Research (22H01627). We thank Hiroki Hosokawa, Dr. Dávid Murányi, and Dr. Yo Miyake for their assistance with the morphology-based identification of macroinvertebrates. Molecular Ecology and Health Laboratory members supported field collection and laboratory work. We thank Mr. Somar Israel Fernando for assisting in the laboratory work. Thanks also to Dr. Naohito Tokunaga of the Division of Analytical BioMedicine, Advanced Research Support Center, Ehime University, for his assistance in performing the Illumina Miseq Sequencing. We thank Dr. Ming-Chih Chiu, Dr. Karen Judan Cruz and Prof. Cristoph Stein-Thoeringer for their valuable comments and suggestions on the earlier version of the manuscript. Thanks to Micanaldo Ernesto Francisco for making the detailed map of the sampling sites.

---

**Supplementary Figure 1.**
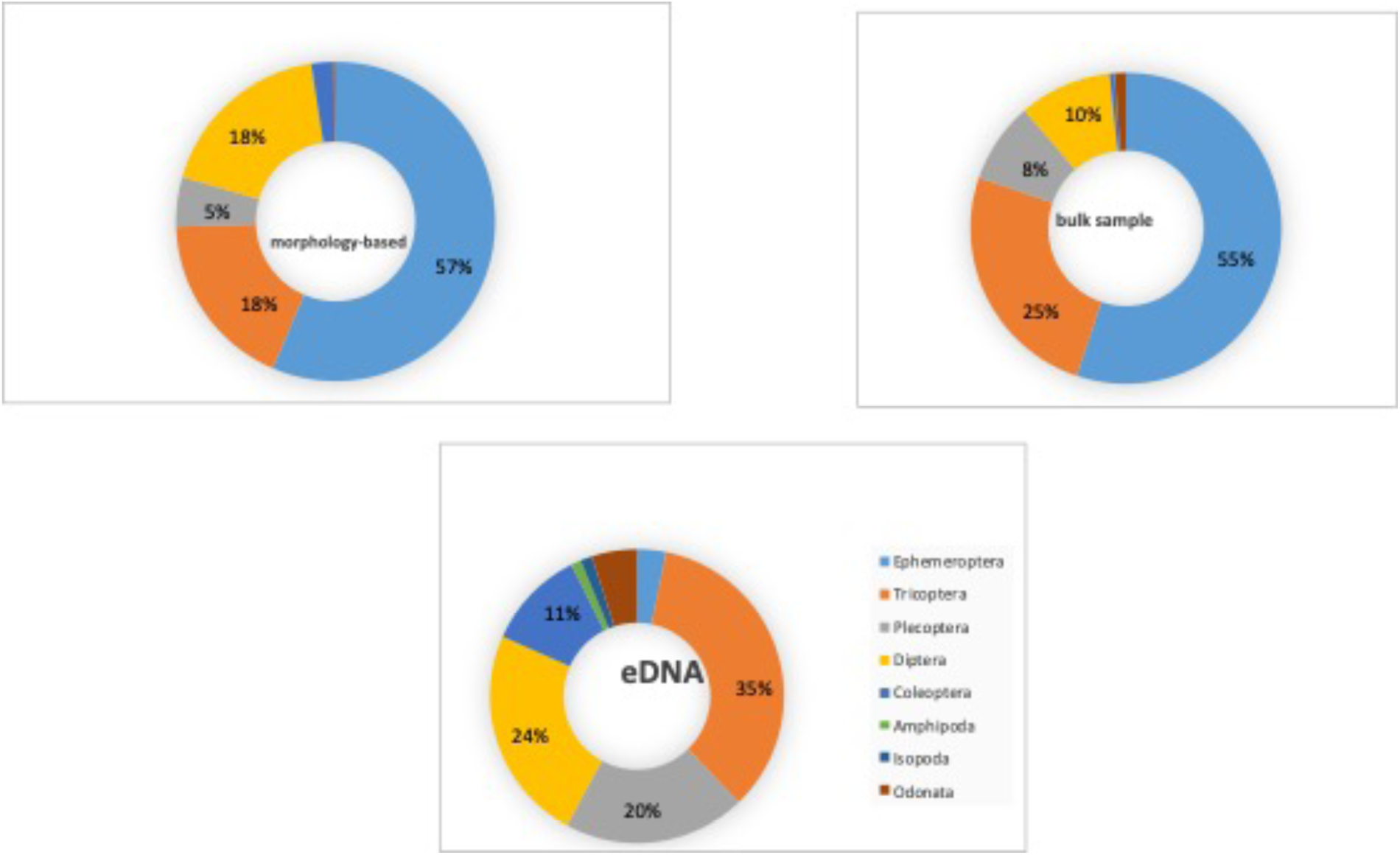
Relative abundabce of taxa for the three approches,e.g., morphology-based identification, bulk-sample, and eDNA metabarcoding based on Phylum.

**Supplementary Figure 2.**
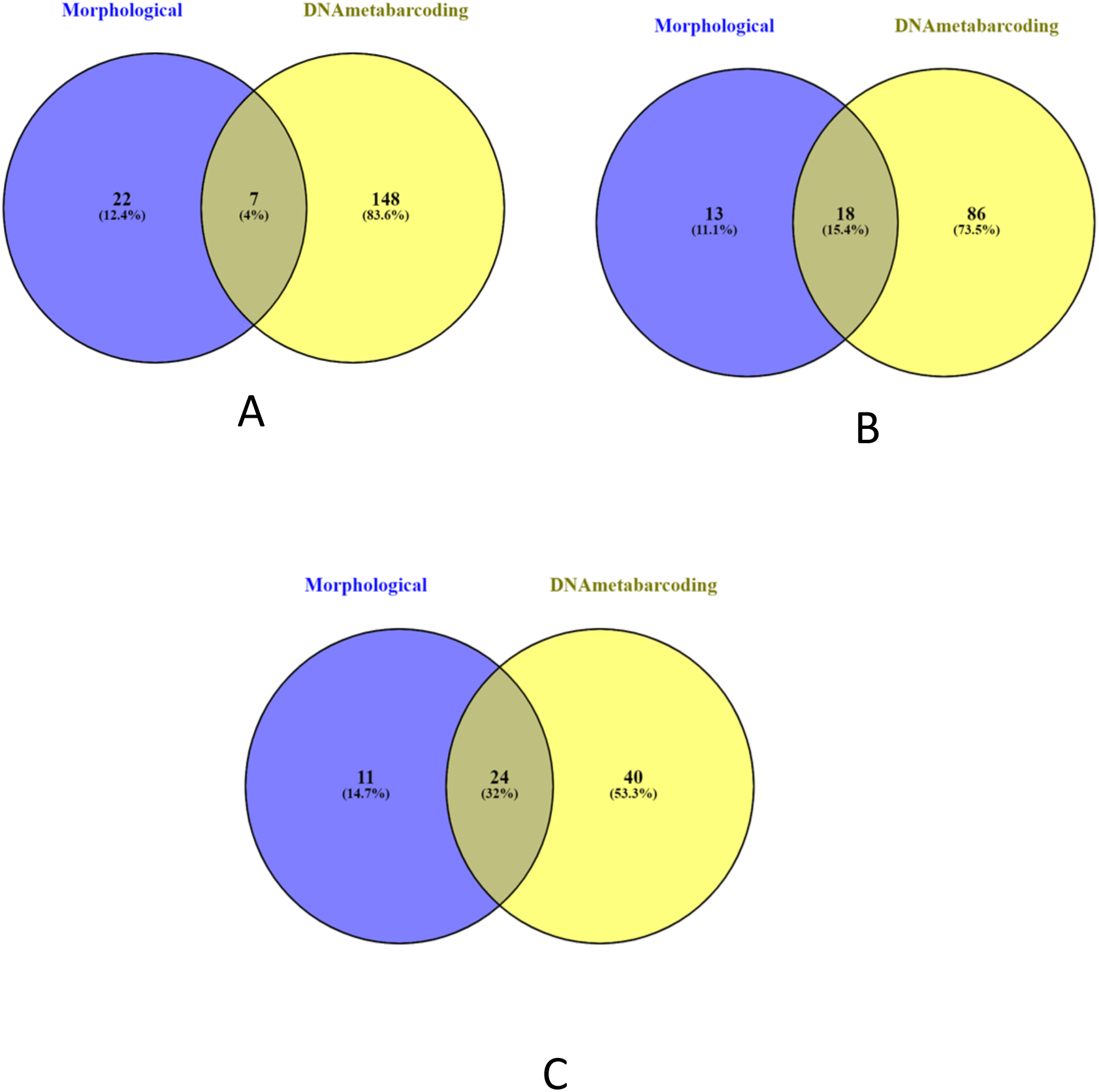
Venn diagrams of taxa sharing by the two approaches, e.g., morphology-based identification approach and DNA metabarcoding (eDNA and bulk-sample) based on the collapsed table (A) Species Level; (B) Genus Level; (C) Family Level.

## References

Andrews, S., (2010). FastQC: A quality control tool for high throughput sequence data.

Aylagas, E. &. R. E., (2014). Environmental status assessment using DNA metabarcoding:towards a genetics-based marine biotic index (gAMBI. PLoS ONE.

Altermatt, F., et al., (2020). Uncovering the complete biodiversity structure in spatial networks: The example of riverine systems. Oikos, 129(5), 607–618.https://doi.org/10.111/oik.06806.

Aylagas, E., et al.,(2018). Adapting metabarcoding-based benthic biomonitoring into routine marine ecological status assessment networks. Ecological Indicators.

Barnes, M. & Turner, C., (2016). The ecology of environmental DNA and implications for conservation genetics. Conservation Genetics, 17(1), pp. 1–17.

Birk, S. et al., (2012). Three hundred ways to assess Europes’s surface waters: an almost complete overview of biological methods to implement the Water Framework Directive. Ecological Indicators, Volume 18, pp. 31–41.

Blackman, R. et al., (2019). Advancing the use of molecular methods for routine freshwater macroinvertebrate biomonitoring-the need for calibration experiments. Metabarcoding and Metagenomics, Volume 3, p. e34735.https://doi.org/10.3897/mbmg.3.34735.

Cahill, A. E., et al., (2018). A comparative analysis of metabarcoding and morphology-based identification of benthic communities across different regional seas. Eco Evol, pp. 8908–8920.https://doi.org/10.1002/ece3.4283.

Callahan, B et al., (2016) DADA2: high-resolution sample inference from illumina amplicon data. Nat. Methods 13, 581–583. Doi:10.1038/nmeth.3869

Carew, M., et al., (2013). Environmental monitoring using next-generation sequencing: rapid identification of macroinvertebrate bioindicator species. Frontiers in Zoology, 10(45), pp. 9–15.

Chao, (1984). Non-parametric estimation of the classes in a population. Scandaivanina Journal of Statistics, 11(4), pp. 265–270.

Chiu, M-C, et al. (2020). Branching networks can have opposing influences on genetic variation in riverine metapopulations. Divers Distrib.; 26: 1813– 1824. https://doi.org/10.1111/ddi.13160.

Cowart, D. et al., (2015). Metabarcoding is powerful yet still blind: a comparative analysis of morphology-based identification and molecular surveys of seagrass communities. PLoS ONE, pp. 10(2),e0117562. 1-26.https://doi.org/10.1371/journal.pone.0117652.

Deiner, K. & Altermatt, F.,(2014). Transport distance of invertebrate environmental DNA in a natural river. PLoS One, 9(2), p. e88786.https://doi.org/10.1371/journal.pone.0088786.

Deiner, K. et al., (2017). Environmental DNA metabarcoding:Transforming how we survey animal and plant communities. Molecular Ecology, Volume 26, pp. 5872–5895.doi:10.1111/mec.14350.

Deiner, K. et al., (2016). Environmental DNA reveals that rivers are conveyer belts of biodiversity information. Nature Communications, Volume 7(1), p. 12544.https://doi.org/10.1038/ncomms12544.

Dudgeon, D., et al. (2006) Freshwater Biodiversity: Importance, Threats, Status and Conservation Challenges. Biological Reviews, 81, 163–182.https://doi.org/10.1017/S1464793105006950

Elbrecht, V. & Leese, F., (2015). Can DNA-based exosystem assessment quantify species abundance? Testing primer bias and biomass-sequence relationships with an innovative metabarcoding protocol. PLoS One, Volume 10, pp. e0130324–16.

Elbrecht, V. & Leese, F., (2016). PrimerMiner:an R package for development and in silico validation of DNA metabarcoding primers. Methods in Ecology and Evolution, pp. 1–5.

Emilson, C. et al., (2017). DNA metabarcoding and morphology-based identification macroinvertebrates metrics reveal the same changes in boreal watersheds across an environmental variable. Scientific Reports, 7(1),12777.https://doi.org/10.1038/s41598-017-13157-x.

Gibson, J. et al., (2014).Simultaneous assessment of the macrobiome and microbiome of bulk-sample of tropical arthropods through DNA metasystematics. PNAS, 11(22), pp. 80007–8012.

Gleason, J. et al., Assessment of stream macroinvertebrate communities with eDNA is not congruent with tissue-based metabarcoding. Molecular Ecology, 30, 3239–3251. https://doi.org/10.111/mec.15597.

Hajibabaei, M. et al., (2019). Watered-down biodiversity? A comparison of metabarcoding results from DNA extracted from matched water and bulk tissue biomonitoring samples. PLoS One, 14(12), p. e0025409.

Hajibabaei, M. et al., (2011). Environmental barcoding: A next-generation sequencing approach for biomonitoring applications using river benthos. PLoS One, 6(4), p. e17497.

Juergens, C., Meyer-Heß, M.F. (2022). Experimental Analysis of Geo-spatial Data to Evaluate Urban Greenspace: A Case Study in Dortmund, Germany. KN J. Cartogr. Geogr. Inf. 72, 153–171. https://doi.org/10.1007/s42489-022-00107-5.

Jørgensen, S.E., et al., (2010). Application of indicators for the assessment of ecosystem health. In S.E. Jørgensen, F. L, Xu & R. Costanza (Eds.,(Handbook of Ecological Indicators for Assessment of Ecosystem Health, 2^nd^ ed.(pp.9–75).CRC Press.

Kawai & Tanida, (2005). Aquatic Insects of Japan: Manual with keys and Illustrations. s.l.:Tokai University Press.

Keck, F et al (2017). Freshwater biomonitoring in the Information Age. Frontiers in Ecology and the Environment, 15(5), 266–274.doi:https://doi.org/10.1002/fee.1490.

Macher, J. & Leese, F.,(2017). Environmental DNA metabarcoding of rivers: Not all eDNA is everywhere, and not all the time available at https://www.biorxiv.org/content/10.1101/164046v1.full.

Mächler, E. et al., (2018). Shedding light on eDNA: Neither natural levels of UV radiation nor the presence of a filter feeder affects eDNA-based detection of aquatic organisms. PloS One, 13(4), p. e0195529.https://doi.org/10.1002/edn3.33.

Martin, M., (2011). Cutadapt removes adapter sequences from high-throughput sequencing reads. Embnet Journal, 17(1), 10–12.

Miyake, Y., et al., (2021). Assessing invertebrate response to an extreme flood event at a regional scale utilizing past survey data. Limnology 22, 169–177 (2021). https://doi.org/10.1007/s10201-021-00651-5

Oksanen, J., et al. (2016). vegan: Community Ecology Package. R package version 2.4-3. Vienna: R Foundation for Statistical Computing.

Patrício, J. et al., (2016). European Marine Biodiversity Networks: Strengths, Weaknesses, Opportunities and Threats. Frontiers in Marine Science, Volume 3.

Pawlowski, J., et al., (2018). The future of biotic indices in the ecogenomic era: Integrating (e)DNA metabarcoding in biological assessment of aquatic ecosystems. Science of The Total Environment, 637-638, 1295– 1310.

R Development Core Team. (2021). “R: A language and environment for statistical computing, reference index version 4.2.1.,” in (Vienna: R Foundation for Statistical Computing). Available at: http://www.R-project.org/.

Rosenberg, D.M., Resh, V.H., (1993). Freshwater biomonitoring and benthic invertebrates. 843 Chapman & Hall, New York, NY

Schenk, N. et al., (2020). Comparison of morphology-based identification, DNA barcoding, and metabarcoding characterization of freshwater nematode. Ecology and Evolution, Volume 10, pp. 2885–2899.

Serrana, J., et al., (2019). Comparison of DNA metabarcoding and morphology-based identification for stream macroinvertebrate biodiversity. Ecological Indicators, Volume 101, pp. 963–972. https://doi.org/10.1016/j.ecolind.2019.02.2008.

Taberlet, A., (2018). Environmental DNA: For Biodiversity Research and Monitoring. Oxford University Press.

Taberlet, P., et al., (2012). Towards next-generation biodiversity assessment using DNA metabarcoding. Molecular Ecology, 21(8), pp. 2045–2050.

Turunen, J. et al., (2021). The power of metabarcoding: Can we improve bioassessment and biodiversity survey of stream macroinvertebrate communities? Metabarcoding and Metagenomics, Volume 5, pp. 99–110.

UNESCO, (1969). Determination of photosynthetic pigments in sea-water. In: Monograph on Ocean Methodology, p. 66.

Valentini, A. et al., (2016). Next-generation monitoring of aquatic biodiversity using environmental DNA metabarcoding. Molecular Ecology, 25(4), pp. 929–942.https://doi.org/10/111/mec.13428.

Weigand, H. et al., (2019). DNA barcode reference libraries for the monitoring of aquatic biota in Europe: Gap-analysis and recommendations for future work. Science of Total Environment, Volume 678, pp. 499–524 https://doi.org/10/1016.j.scitotenv.2019.04.247.

Yates, M. et al. (2021). Integrating physiology and environmental dynamics to operationalize environmental DNA(eDNA) as a means to monitor freshwater macroorganisms abundance. Molecular Ecology, Volume 30, pp. 6531–6550.

Zhou, X. et al., (2013). Ultra-deep sequencing enables high-fidelity recovery of biodiversity for bulk arthropod samples without PCR amplification. Giga Science, 2(4).

Zimmerman, J. et al., (2015). Metabarcoding vs. morphology-based identification to assess diatom diversity in environmental studies. Molecular Ecology, Volume 15, pp. 526–542. https://doi.org/10.1111/1755-0998.12336.

